# How does host age and nutrition affect density regulation of obligate versus facultative bacterial symbionts? Insights from the tsetse fly

**DOI:** 10.1101/2024.09.13.612807

**Authors:** Mathilda Whittle, Antoine M.G. Barreaux, Lee R. Haines, Michael B. Bonsall, Sinead English, Fleur Ponton

**Affiliations:** Faculty of Life Sciences, University of Bristol, Bristol, UK; School of Natural Sciences, Macquarie University, NSW, Australia; UMR INTERTRYP, CIRAD, Montpellier, France; Animal Health Theme, ICIPE, Nairobi, Kenya; Department of Biological Sciences, University of Notre Dame, Indiana, USA; Department of Biology, University of Oxford, Oxford, UK; St Peter’s College, Oxford, UK

## Abstract

The relationships between insect hosts and their symbionts can vary tremendously in the extent to which hosts depend on and control their symbionts. Obligate symbionts that provide micronutrients to their host are often compartmentalised to specialised host organs and depend on their hosts for survival, whereas facultative symbionts retain the ability to survive outside of their hosts. Few studies compare the extent to which a host controls and adjusts the density of obligate and facultative symbionts directly. Here, we used tsetse as a model for teasing apart the relationships between a host (*Glossina morsitans morsitans*) and obligate (*Wigglesworthia glossinidia*) and facultative (*Sodalis glossinidius*) symbionts. We hypothesised that tsetse actively regulate the density of *Wigglesworthia* according to the host’s requirements, depending on their current nutritional state and developmental age. In contrast, we postulated that *Sodalis* retains some independence from host control, and that the growth of this symbiont is dependent on the conditions of the immediate environment, such as nutrient availability. Using qPCR, we examined how symbiont densities change across host age and the hunger cycle. Additionally, we investigated how host nutrition influences symbiont density, by comparing tsetse that were fed diluted blood (poor nutrition) or blood supplemented with yeast extract (vitamin enriched). We found that the density of *Wigglesworthia* did not reflect the nutritional status of the host, but was optimised to accommodate long-term host requirements (in terms of nutrient provisioning). In contrast, the density of facultative *Sodalis* was influenced by the ecological context (i.e. nutrient availability). This suggests that tsetse regulate the abundance of *Wigglesworthia* to a greater extent than *Sodalis*. We propose that tsetse exert only partial control over *Sodalis* growth due to the relatively recent transition of this symbiont to host-associated living.

**Author summary:** Symbiotic microbes have the potential to significantly impact the wider ecosystem by affecting the fitness and behaviour of their animal hosts. The density of a particular symbiont population within host tissues is likely an important factor influencing the effect it has on the host, however, little is known about the factors which determine how symbiont density is regulated, and how these differ between symbionts with different degrees of host-association (e.g. obligate and facultative symbionts). Here, we found that *Wigglesworthia* and *Sodalis*, two bacterial tsetse symbionts, demonstrate distinct trends in density according to host age and nutrition. We discuss how the evolutionary histories of these symbionts with their host potentially explain these results, highlighting the complexity and dynamic nature of host-symbiont interactions. Our findings contribute to our understanding of the extent to which hosts and symbionts control symbiont density and how symbiont density regulation can be affected by the ecological context.

## Introduction

Many eukaryotic organisms, such as animals, plants and protists, live with symbiotic microorganisms within their bodies (1-3). A great diversity exists in the nature of such symbioses. On one end of the spectrum, parasitic symbionts exploit the host for their own benefit, while on the other end, mutualistic symbionts provide benefits that increase host fitness (4). There are many examples of mutualistic associations between microbes and hosts, such as photosynthetic algae providing essential nutrients to cnidarian hosts (5), protective bacteria in nematodes producing chemicals to defend their host from pathogens (6), and symbionts improving their insect hosts’ tolerance to heat stress (7). Some mutualisms can evolve to become obligate for both the host and the symbiont, i.e., the host depends on the symbiont for survival and vice versa (8).

Supporting a symbiont population, even an obligate one, entails a metabolic cost for the host because symbionts acquire all their nutritional resources from within the host (9). The net benefit (i.e. the difference between the benefit provided to the host and the cost of supporting the symbiont) for a host participating in a mutualistic symbiosis depends on the ecological context (10-17). For instance, the diet of many insects is supplemented with micronutrients produced by gut symbionts (18, 19), and the associated benefit depends on the host’s nutritional requirements and the availability of such micronutrients in the host’s diet. As the size of the symbiont population within the host tissues likely correlates with the amount of nutrients provisioned to the host, as well as the metabolic cost of maintaining the symbiont, active host regulation of symbiont densities according to the host requirements and the availability of dietary nutrients could allow hosts to maximise the net benefits (20). Empirical studies suggest that symbiont density regulation according to host requirements occurs in several obligate nutritional symbioses of insects (reviewed in 21). Several examples of this are: aphids harbour different symbiont densities according to their host plant (13); weevils remove their symbiont after maturation (16); and female tsetse flies (which have comparatively large reproductive investment (22)) host higher symbiont densities than the males (11, 23).

Many obligate symbioses of insects are ancient associations (24). Long coevolutionary histories between hosts and symbionts have resulted in highly reduced symbiont genomes, which limit the ability of symbionts to survive outside hosts and regulate their own replication (25). Additionally, several mechanisms that provide hosts with control over symbioses have been identified (21). In particular, the evolution of specialised symbiont-housing organs has occurred independently in many hosts, and facilitates control over the localisation and abundance of their obligate symbionts (26). In contrast to obligate symbionts, facultative symbionts, although potentially beneficial, are not strictly required by the host for survival, and often represent more recent transitions to host-associated living (27). Many facultative symbionts have retained the ability to survive outside of host tissues, likely due to lower gene erosion, and demonstrate a more extensive tissue distribution in the host body (28).

Here, we investigate the degree of control simultaneously exerted by a host over the density of an obligate and a facultative symbiont. One issue of using symbiont density to infer host control of a symbiosis lies in attributing changes in density to host-mediated regulation (as opposed to any direct effects of the ecological context on bacteria growth). By comparing two symbionts of the same host with different evolutionary histories, under several ecological scenarios, we aimed to provide a broad set of evidence for the potential influence of host control on symbiont density. We hypothesised that while the host may maintain and adjust the density of its obligate symbiont according to its requirements, the facultative symbiont retains a greater degree of independence from the control of its host. As such, growth of the facultative symbiont is likely to be more directly affected by the immediate environment, i.e. the nutrient availability within host tissues.

We tested our hypotheses using tsetse (*Glossina* spp.), and their association with obligate (*Wigglesworthia glossinidia*) and facultative (*Sodalis glossinidius*) symbionts, as a model system. Tsetse are exclusive blood feeders, and rely on *Wigglesworthia* for crucial B vitamins not available in sufficient concentrations in their diet (29). Having co-evolved with tsetse for 50–80 Ma (30), *Wigglesworthia* has achieved a highly integrated role in tsetse biology as characterised by an extremely reduced genome (0.7 Mb)(31, 32) and mutually obligate status with all *Glossina* hosts (30, 33). The primary population of *Wigglesworthia* is located intracellularly, within the specialised organ known as the bacteriome (mycetome), which saddles the anterior midgut. Female tsetse, unlike most flies, do not lay eggs but produce one offspring at a time. The single larva is fed on a milk-like substance *in utero* (34), and a secondary population of *Wigglesworthia* exists within the milk glands, which allows the maternal transmission of this symbiont to tsetse offspring (35). Compartmentalisation of symbionts to specialised housing organs, such as the bacteriome or milk glands, helps a host control its symbiont populations both in maternal and offspring generations (26), and as such, tsetse may have the ability to tightly regulate the abundance of *Wigglesworthia*.

The tsetse secondary bacterial symbiont, *Sodalis glossinidius*, is widespread throughout insectary and field tsetse populations and represents a relatively recent transition from free-living bacteria to an endosymbiotic lifestyle (36). Accordingly, *Sodalis* can colonise multiple tsetse tissues including the hemolymph, midgut, fat body, milk glands (in females, where it is transmitted to host offspring alongside *Wigglesworthia* (35)), and testes and spermatophore (in males, where it can be transmitted horizontally during mating (37)). Within tsetse tissues, *Sodalis* demonstrates a dependency on thiamine (vitamin B_1_) produced by *Wigglesworthia* as it lacks the ability to produce this vitamin itself (38). However, *Sodalis* retains the ability to survive and grow outside of host tissues in *in vitro* cultures (39). How tsetse benefit from *Sodalis* is uncertain; although the bacteria is not found in all tsetse populations, antibiotic removal of *Sodalis* from tsetse appears to result in decreased longevity and reduced susceptibility of the host to trypanosome infection (40).

Here, we used qPCR to measure the density of *Wigglesworthia* and *Sodalis* associated with the digestive and reproductive tissues of tsetse, under several ecological scenarios. First, we measured symbiont density at multiple time points across adult female age. Following an initial proliferation in early adulthood (as is observed of symbionts of several species, including tsetse (10, 17, 41-45)), we predicted that *Wigglesworthia* may be maintained at a constant level to support the nutritional demands of reproduction (see Fig 1a for an illustration of the directional predictions). As thiamine provisioning from *Wigglesworthia* increases, we predicted that the density of *Sodalis* would increase in parallel to *Wigglesworthia* (17). Additionally, maturation and senescence of the host immune system (46, 47) may reduce the ability of tsetse to control the density of *Sodalis* in early adulthood and ageing hosts, respectively, thus proliferation may be observed at these periods (Fig 1a). Second, we measured symbiont density in tsetse following a blood meal. Proliferation of gut microbiota following a blood meal has been observed in mosquitoes (48, 49), therefore we predicted that *Sodalis* may demonstrate brief proliferation in the nutrient-rich environment, but tsetse will maintain a constant density of *Wigglesworthia* (Fig 1b) according to the host’s long-term requirements for B vitamins. Third, we measured symbiont density in tsetse receiving nutritionally manipulated diets: diluted blood with reduced nutritional content, or blood supplemented with yeast extract thereby enriched with B vitamins (as in 50). Where energy is limiting, supporting a symbiont population could become relatively more costly to the host, and so we predicted that the density of *Wigglesworthia* will be lower in hosts receiving nutritionally-poor blood (Fig 1c). *Sodalis* may also grow to lower densities in hosts receiving poorer diets (Fig 1c) due to fewer resources being available within host tissues. Providing an exogenous source of B vitamins undermines the benefit *Wigglesworthia* provides to tsetse, therefore we predicted that the density of this symbiont will be actively reduced by hosts receiving an enriched diet as it is surplus to requirements (Fig 1d). In contrast, we predicted that *Sodalis* may proliferate in the presence of a nutritionally-enriched environment (i.e. containing macronutrients and thiamine from yeast extract) (Fig 1d).

**Fig 1.**
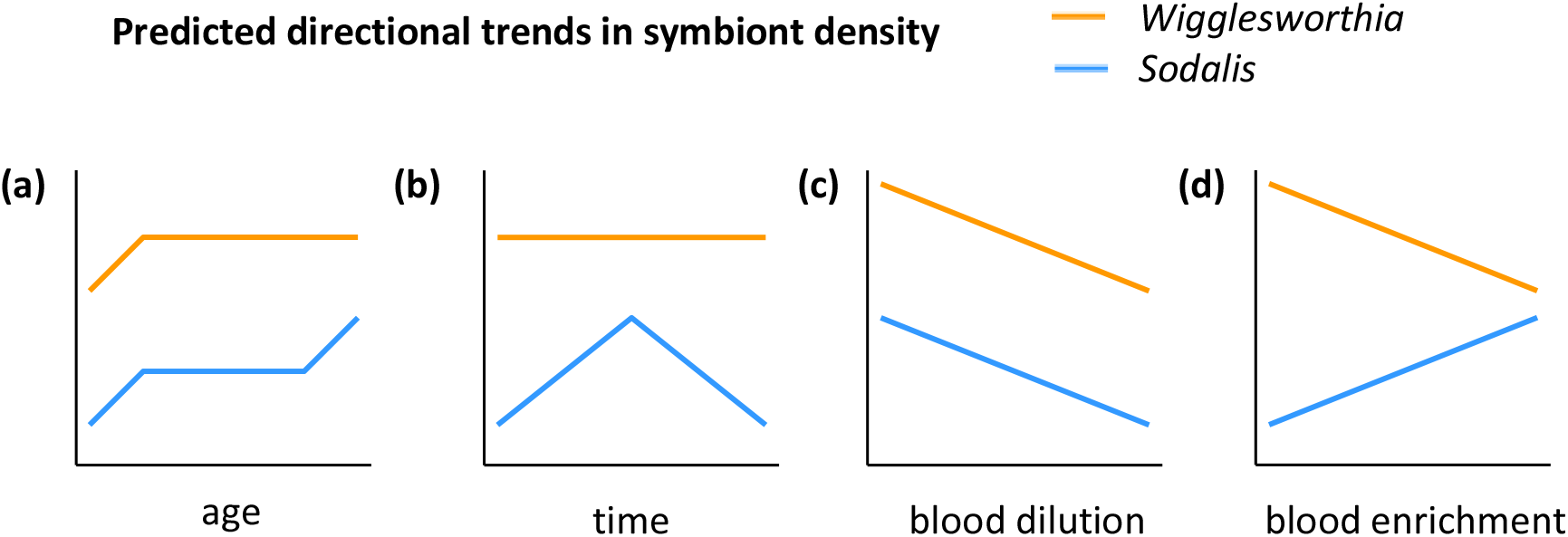
Predicted directional trends in the density of *Wigglesworthia* and *Sodalis*: (a) Study 1: across adult female age, (b) Study 2: over 96 hours following a blood meal, (c) Study 3: from hosts reared on diets of diluted blood, and (d) Study 3: from hosts reared on diets of blood enriched with yeast extract. The orange and blue lines indicate trends in symbiont density rather than the densities of *Wigglesworthia* and *Sodalis* relative to each other.

## Materials and Methods

### Tsetse rearing

We obtained *Glossina morsitans morsitans* from the Liverpool School of Tropical Medicine – either at the adult stage (study 1: female age) or as pupae (study 2: hunger cycle, and study 3: dietary manipulation experiments). Individuals were maintained at 25 °C with 70% humidity, with 12:12 hours light: dark cycles. Flies were fed on sterile defibrinated horse blood (TCS Biosciences, UK) three times a week (Monday, Wednesday and Friday) using an artificial membrane system.

### Study 1: symbiont density dynamics across female age

For investigating symbiont density dynamics across adult age, we used females from 16 different age groups spanning the age range of the tsetse colony; from approximately 6 hours to 88 days post emergence, with n = 10 for each group. Flies were killed by freezing at -80 °C, 48 hours following their final blood meal (excluding newly emerged flies, which were unfed) and processed for qPCR quantification of symbiont density (see below). Upon maturation, female tsetse produce offspring at regular intervals (every 9-10 days) throughout adulthood. Within individuals, the density of *Wigglesworthia* and *Sodalis* is likely to fluctuate during adulthood due to transmission of both symbionts to developing larvae (51), however, we anticipated that these short-term fluctuations were not likely to be observed at a population level due to the reproductive cycles being unaligned between females.

### Study 2: symbiont density depending on hunger levels in males

We reared newly emerged males for approximately 4 weeks on a standard horse blood diet. We anticipated that by using males, small changes in symbiont density across the hunger cycle may be observed, as any contribution of fluctuating symbiont populations in the female milk glands are eliminated. We caged males without females to prevent any reduction in *Sodalis* density due to mating (37). To maximise likelihood of feeding and to allow complete digestion of the previous meal, flies were starved for 4 days prior to the final meal (at 4 weeks old). All flies appeared fully engorged at the final meal. Flies were killed by freezing at -80 °C at 6 (n = 13), 24 (n = 14), 48 (n = 12) or 96 (n = 13) hours following the final meal and processed for qPCR quantification of symbiont density.

### Study 3: effect of dietary manipulation on symbiont density in females

Adult females were caged upon emergence at a female: male ratio of 4:1. We reared flies for 6 weeks on one of 10 dietary treatments (table 1). We used normal saline (0.9% NaCl (w/v) in water) to reduce the nutritional content of the diet at varying ratios, and yeast extract to fortify the diet with B vitamins at increasing concentrations. For addition to the blood diet, we dissolved 0.4, 0.8, 1.6 and 4 g of dried yeast extract (Merck, cat. No. 70161) in aliquots of 10 mL reverse osmosis (RO) water, which we then sterilised using syringe filters (0.2 μm). The yeast solutions were then added to aliquots of 70 mL blood to create blood enriched with 0.5, 1, 2 and 5% w/v yeast extract.

**Table 1.** Treatment groups for dietary manipulation (study 3). Flies were reared for six weeks (from emergence) on their respective treatments. Surviving flies were processed for qPCR analysis.

| Group | Treatment | Sample size at start | Sample size for analysis |
| --- | --- | --- | --- |
| Control | Standard defibrinated horse blood | 15 | 12 |
| Blood diluted with normal saline | 3:1 v/v blood: saline | 15 | 13 |
|  | 2:1 v/v blood: saline | 15 | 13 |
|  | 1:1 v/v blood: saline | 15 | 10 |
|  | 0.5:1 v/v blood: saline | 15 | 12 |
|  | 0.3:1 v/v blood: saline | 15 | 8 |
| Blood supplemented with yeast extract | 0.5% w/v yeast extract | 10 | 9 |
|  | 1% w/v yeast extract | 10 | 9 |
|  | 2% w/v yeast extract | 10 | 9 |
|  | 5% w/v yeast extract | 10 | 3 |

Diets were pre-prepared with fresh blood and stored for up to two weeks at 3°C. To ascertain any detrimental effects of the treatments on host survival, we counted the number of living females daily. Pupae were collected from each group and incubated under the same environmental conditions until emergence. We recorded the weight at deposition of pupae produced by each treatment group, and the number of days for the unfed adult offspring to die post emergence, to determine if the dietary treatments affected host reproduction and resource provisioning to offspring. All surviving females were killed by freezing at -80°C, 48 hours after their final blood meal, and processed for qPCR quantification of symbiont density.

### Sample preparation and DNA extraction

We measured the length of the “hatchet” wing cell as a standard measure for host size (52). The total content of the abdomen was removed by dissection into 180 μL ATL buffer from the DNeasy blood and tissue kit (Qiagen, CA). Instruments and surfaces were washed with 70% ethanol to prevent cross-contamination. We estimated the stage of pregnancy (i.e. egg or 1^st^, 2^nd^ or 3^rd^ larval instar) for females (53), and removed any 2^nd^ and 3^rd^ instar larvae from the samples. Total genomic DNA of the remaining abdominal content was extracted using the DNeasy blood and tissue kit (Qiagen, CA), following a modified version of the manufacturer’s protocol (supplementary material). Extracted DNA was desiccated for short-term storage at -30°C using an Eppendorf Vacufuge plus (Eppendorf, UK). Dried DNA was then rehydrated with 50 µL PCR grade water (Fisher Biotec, Australia) for analysis.

### Primers

We used pairs of specific primers (for PCR and qPCR amplification) for species-specific and single-copy genes of *G. morsitans morsitans* (*alpha-tubulin*, GenBank Accession no. ADD19945.1), *Wigglesworthia glossinidia* (thiamine biosynthesis protein (*thiC*), GenBank Accession no. AGG38086.1) and *Sodalis glossinidius* (*exochitinase*, GenBank accession no. BAE74749.1) (table S1).

### Assay optimisation and standard curves

We used non-experimental flies to obtain PCR products by amplification of *alpha-tubulin, thiC* and *exochitinase* genes (see supplementary material for cycling conditions), which we then purified using QIAquick PCR purification kit (Qiagen, CA). DNA concentration was measured using a Qubit 4 fluorometer (Invitrogen, MA). To determine the linear dynamic range for each target gene, we constructed standard curves using four-fold serial dilutions (over nine points) of PCR products of known concentration (concentration ranges given in table S2). We verified the efficiency of amplification and dynamic range for purified PCR products (91.8-97.0%) against standard curves constructed from total DNA from non-experimental flies (95.4-98.3%). We confirmed the specificity of primer annealing by performing a melt-curve analysis.

Quantitative (Real Time) PCR amplification was performed using a CFX384 Touch Real-Time PCR Detection System (Bio-Rad, CA) for each primer pair (see supplementary material for cycling conditions). Amplifications were obtained using PowerUP SYBR Green master mix (Applied Biosystems, MA) in a final volume of 10 µL, including 4 µL of template. We found optimal primer concentrations by running titrations between 250 nM and 900 nM of the Forward and Reverse primers. Primer concentration of 500 nM was chosen as it yielded the lowest quantification cycle (C_q_) for all primer pairs. We performed each assay in triplicate, including no template controls.

### Calculation of symbiont abundance

We diluted the DNA samples with PCR grade water (Fisher Biotec, Australia) to fit within the dynamic range of the standard curves (table S2). We included standard curves on each qPCR plate for absolute quantification of the target genes (table S2). The C_q_ for each sample was used to estimate the quantity of target DNA (in nanograms) initially present in the template by comparing against the standard curve. We then calculated the number of gene copies according to equation 1 (45), where the amplicon size is given by the number of base pairs (table S1). As symbiotic bacteria, including *Wigglesworthia*, can demonstrate polyploidy (17), we defined the symbiont density as the number of symbiont genomes divided by the number of *G. morsitans* genomes.

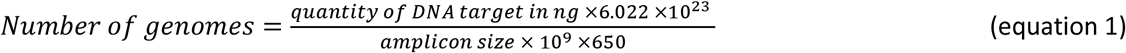

### Statistical analysis

We analysed data using R (version 4.2.1) (54). Density data were log transformed to satisfy normality and analysed using multiple linear regression. To elucidate the functional form of trends in *Wigglesworthia* and *Sodalis* density, we used Akaike’s information criterion (corrected for small sample size, AICc) model selection, to compare models with increasing order polynomials (see below for each analysis). Models with the lowest AICc were selected as the best-fit and compared to our *a priori* predictions for each symbiont. The regression models visualised in Figs 2-4 were created using the best-fit models. The difference in AICc between each model and the selected model (ΔAICc), and the Akaike’s weights (ω_i_), were calculated for each model (55). The Akaike’s weights are interpreted as the probability that each model is the best model of the set, in terms of AICc (56). The length of the hatchet wing cell (all flies) and stage of pregnancy (females only) were included as fixed effects in all relevant models. Model assumptions were verified using the “performance” package (57).

**Fig 2.**
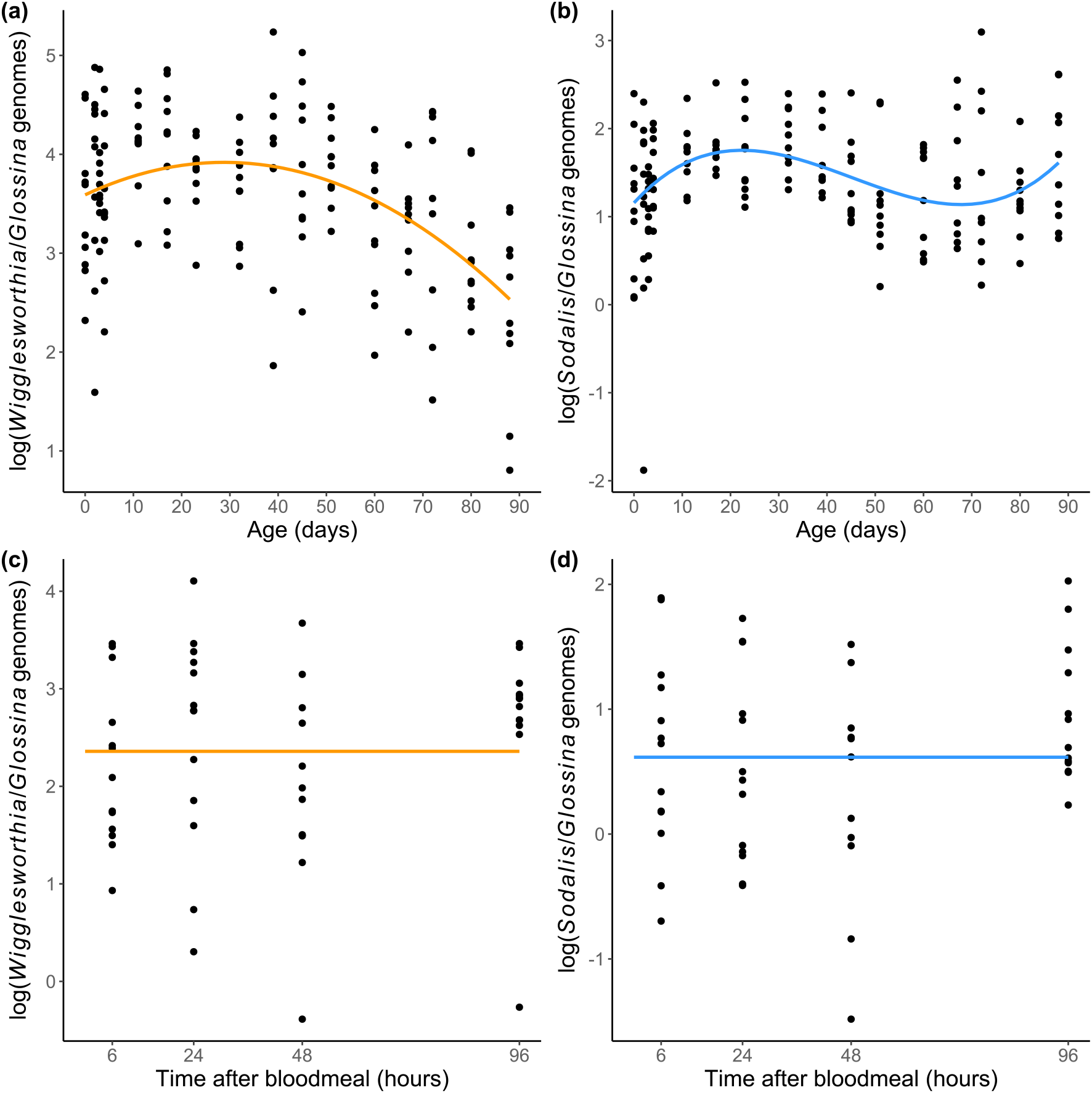
Density dynamics of *Wigglesworthia* (orange) and *Sodalis* (blue) (a-b) throughout adult development (females), and (c-d) throughout the hunger cycle (4-week old males). Regression lines indicate predictor effects of selected models (tables S3-S4). Note that the y-axis is on different scales between the two symbionts.

**Fig 3.**
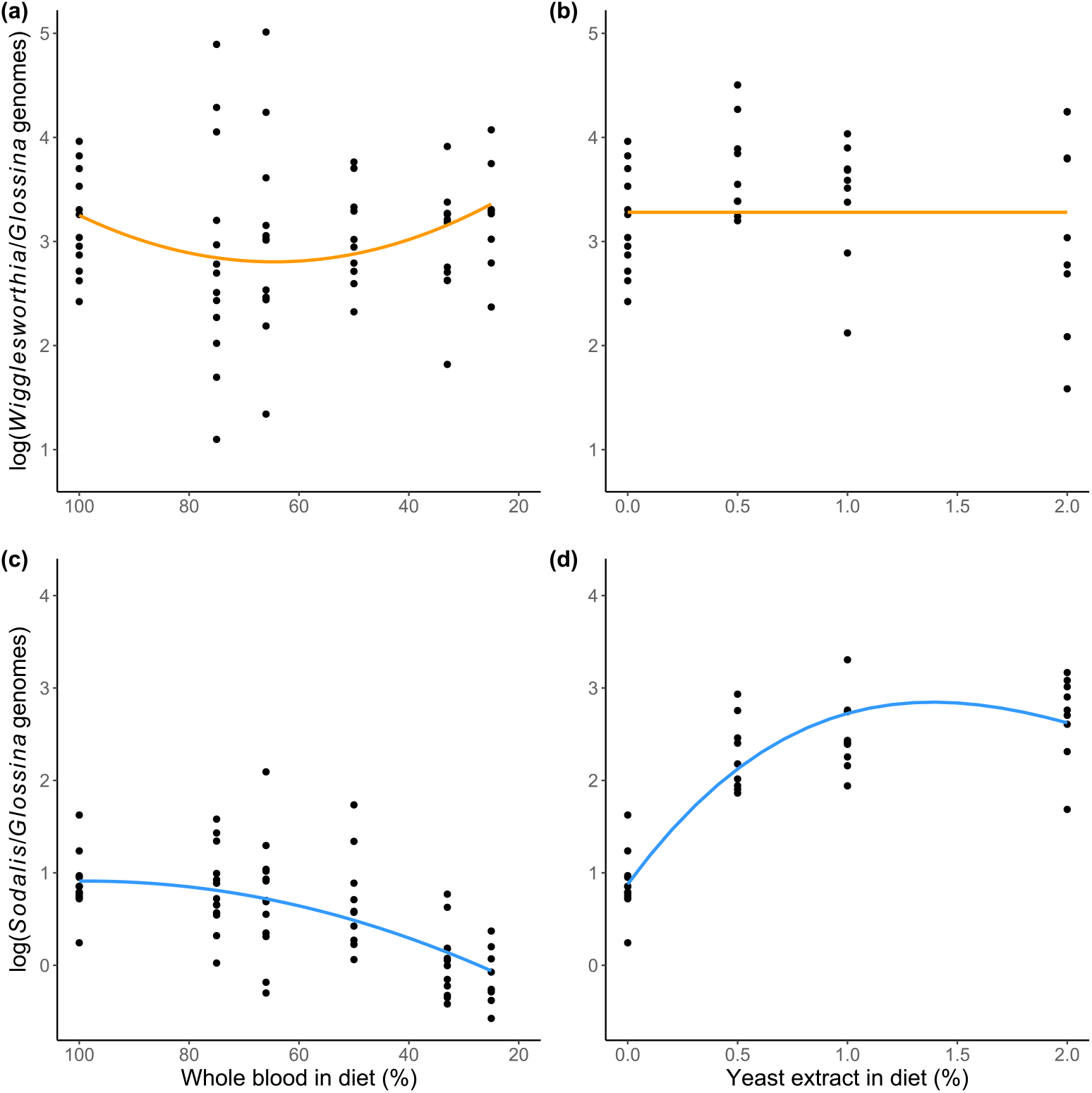
Effect of dietary manipulation on the density of *Wigglesworthia* (orange) and *Sodalis* (blue) in 6-week old female tsetse. (a) *Wigglesworthia* density according to blood concentration (%). (b) *Wigglesworthia* density according to yeast extract (% w/v). (c) *Sodalis* density according to blood concentration (%). (d) *Sodalis* density upon yeast extract (% w/v) supplementation. Blue lines indicate predictor effects of selected models (table S5). Note that the y-axis is on different scales between the two symbionts.

**Fig 4.**
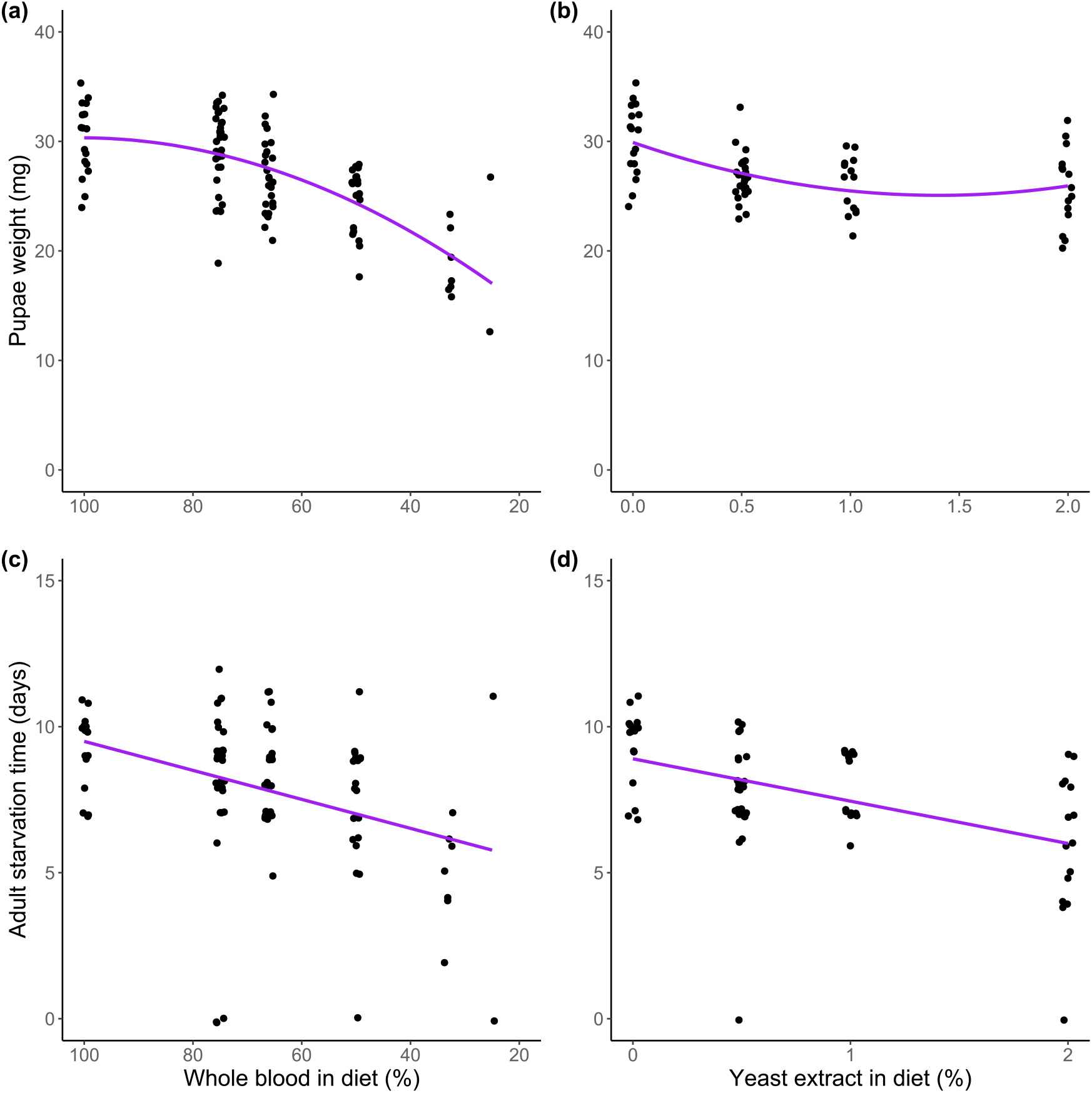
Effect of blood dilution with saline and blood enrichment with yeast extract on host reproductive output. (a-b) Weight of pupae at time of deposition. (c-d) Time taken for unfed offspring to starve upon emergence. Purple lines indicate predictor effects of selected models (table S6).

### Study 1: symbiont density dynamics across female age

To compare the dynamics of *Wigglesworthia* and *Sodalis* across female age, we fit models including intercept-only, linear, quadratic and cubic effects of host age (in days), then selected the model with the lowest AICc for each symbiont.

### Study 2: symbiont density depending on hunger levels in males

To compare the dynamics of *Wigglesworthia* and *Sodalis* throughout the hunger cycle, we fit models including intercept-only, linear, quadratic and cubic effects of the amount of time since feeding (in hours), then selected the model with the lowest AICc for each symbiont.

### Study 3: effect of dietary manipulation on symbiont density in females

The effects of rearing hosts on blood diluted with saline and blood enriched with yeast extract were analysed separately. To analyse the effect of blood content in diet, we calculated the percentage of whole blood in each treatment for use as a continuous predictor. The effect of blood dilution on *Wigglesworthia* and *Sodalis* densities was investigated by fitting models including intercept-only, linear, quadratic and cubic effects of the amount of defibrinated blood in host diet (%), then the model with the lowest AICc was selected for each symbiont. Likewise, the effect of blood enrichment on *Wigglesworthia* and *Sodalis* densities was investigated by fitting models including intercept-only, linear, quadratic and cubic effects of the amount of yeast extract in host diet (% w/v), inputted as a continuous variable. The same group of flies receiving standard horse blood (i.e. 100% defibrinated blood and 0% yeast extract) were used in both analyses as controls. Due to very high tsetse mortality rates when fed blood containing 5% yeast extract (resulting in a small sample size, Fig S1), we excluded this group from the analysis.

The effects of blood content and blood enrichment on host reproduction were found using the same model selection methodology as above. We fit models including intercept-only, linear, quadratic and cubic effects of the amount of whole blood in host diet (%) on pupae weight (in mg) and survival time post emergence (in days), then selected the models with the lowest AICc for each measure of success. Likewise, we fit models including intercept-only, linear, quadratic and cubic effects of the amount of yeast extract in host diet (%) on pupae weight and starvation time upon emergence, then selected the models with the lowest AICc. The most diluted blood treatment (1:3 v/v blood:saline) and most enriched (5% yeast extract) were excluded from the analyses due to the very low reproductive output of hosts reared on these extreme treatments (two and one pupae, respectively).

## Results

### Study 1: symbiont density dynamics across female age

Across the range of adult female ages, the density of *Wigglesworthia* was generally greater than that of *Sodalis* (Fig 2a,b). *Wigglesworthia* proliferated in early adulthood, up to approximately 17 days post-emergence, after which the density of this symbiont plateaued and declined (Fig 2a). Accordingly, we found evidence of a quadratic effect of host age on *Wigglesworthia* density, as determined by the best-fit model carrying 75% of the AIC weight (i.e. ω_i_ = 0.75, table S3). For *Sodalis*, after an initial increase in hosts up to 17 days post-emergence, the density declined, however, this decline did not continue throughout adulthood and increased slightly in hosts from the age of approximately 67 days old (Fig 2b). The model which included the cubic effect of host age was selected as the best-fit for predicting the dynamics of *Sodalis* density and carried 97% of the AIC weight (table S3).

### Study 2: symbiont density depending on hunger levels in males

We found no evidence that the density of either *Wigglesworthia* or *Sodalis* varied depending on time since feeding, indicated by selection of the intercept-only model as the best-fit for both symbionts (Fig 2c,d, table S4).

### Study 3: effect of dietary manipulation on symbiont density and host reproduction in females

The model including a quadratic effect of blood content in diet on *Wigglesworthia* density was determined to be the best-fit, indicating that the density initially decreases with blood dilution and then increases again in hosts receiving very diluted blood (Fig 3a), however, this model only carried 42% of the AIC weight (table S5). The intercept-only model carried 36% of the AIC weight, indicating that the evidence for an effect of blood content on symbiont density is weak. Similarly, we found no evidence that enrichment of blood with yeast extract affects the density of *Wigglesworthia* (Fig 3b), given by selection of the intercept-only model carrying 71% of the AIC weight (table S5). The density of *Sodalis* appeared to decrease slightly with increasingly diluted diets (Fig 3c). Collectively, the linear, quadratic and cubic models carried > 99.9% of the AIC weight, indicating that there is strong evidence that blood content affects the density of *Sodalis*. The density of *Sodalis* appeared to increase substantially when yeast extract was added to the host diet, although this appeared to plateau with increasing enrichment (Fig 3d). Accordingly, we found strong evidence of a cubic effect of blood enrichment on the density of *Sodalis*, given by selection of the cubic model carrying >99% of the AIC weight (table S5).

Both dilution of the blood with saline and enrichment with yeast extract appeared to negatively impact female reproduction in terms of reducing pupae weight and the survival time of adult offspring upon emergence (Fig 4). The detrimental effect of blood dilution on pupae weight appeared to increase with very low blood content in the host diet (Fig 4a), which is moderately supported by selection of the model including a quadratic effect of blood content, carrying 67% of the AIC weight (table S6). Conversely, we found that pupae weight decreased with addition of any yeast extract to host diet, but plateaued with increasing enrichment (Fig 4b). We found moderate evidence for this non-linear effect, given by selection of the model including a quadratic term, carrying 64% of the AIC weight (table S6). We found strong evidence that both blood dilution (Fig 4c) and blood enrichment (Fig 4d) reduced the time taken for adult offspring to starve upon emergence, as both intercept-only models carried less than 0.01% of the AIC weight (table S6).

## Discussion

Here, we investigated the different regulatory forces on the density of *Wigglesworthia* and *Sodalis*, obligate and facultative symbionts of tsetse respectively, within the digestive and reproductive tissues of tsetse. By measuring the density of *Wigglesworthia* and *Sodalis* in several ecological scenarios, we aimed to test predictions of how symbiont density would change with host requirements (e.g. across age, over the hunger cycle and depending on the diet). Moreover, we were interested in whether any patterns differed between the two types of symbionts.

First, we measured symbiont density across females in different age groups. We found that both *Wigglesworthia* and *Sodalis* proliferate following emergence, (Fig 2a,b), which supports previous findings (17, 45). Tsetse generally produce their first offspring around the time that symbionts reach a stable density (approximately 18-23 days (58, 59)). Females may initially invest a large amount of resources into growing the population of *Wigglesworthia* until it is sufficient to meet the B vitamin demands of producing offspring. Upon the onset of reproduction, limiting the resources made available to *Wigglesworthia* and transmission of *Wigglesworthia* to offspring at regular intervals may prevent the continued growth of the *Wigglesworthia* population (51). Maturation of the immune system in teneral adults is known to leave them susceptible to infection (46), and this may also give an opportunity for symbiont proliferation during this period. Compartmentalisation of symbionts to bacteriomes serves to protect symbionts from immune effectors circulating in the haemolymph (60), so a weakened immune function of the host may have little effect on the primary population of *Wigglesworthia* density. It is possible, however, that regulation via the immune system is a mechanism for hosts to limit *Sodalis* density. The concordant proliferation of *Wigglesworthia* and *Sodalis*, which plateaus at similar times (approximately 17 days, Fig 2a,b), emphasises the between-symbiont interactions that influences *Sodalis* density. Where dependency on nutrients (e.g. thiamine (38)) produced by the obligate symbiont limits the proliferation of the facultative symbiont, the host may not need to invest greatly in controlling the abundance of *Sodalis* itself.

We hypothesised that due to constant reproduction throughout adult development in female tsetse, the demands for B vitamins would be constant and therefore the density of *Wigglesworthia* would be maintained at a constant level. We found that after the initial proliferation, both *Wigglesworthia* and *Sodalis* appeared to decrease in density in digestive and reproductive tissues (Fig 2a,b), contrary to our predictions (Fig 1a). Previous investigations of symbiont abundance in mated adult female tsetse found no positive or negative trend in *Wigglesworthia* during adulthood (17, 45), however, these findings were based on females of up to 8 weeks (56 days) old, and our observed decrease in *Wigglesworthia* is most noticeable in flies older than 8 weeks (Fig 2a). Studies of the aphid-*Buchnera* system, a nutritional symbiosis analogous to tsetse-*Wigglesworthia*, observed degradation of the bacteriocytes (the cells which contain the symbiotic bacteria and form the bacteriome) in senescent hosts (61), and that the number of genomes per *Buchnera* cell decreased in old hosts (62). Similar effects of ageing in tsetse may explain the observed decline in *Wigglesworthia* density (Fig 2a). Previous studies quantifying *Wigglesworthia* in adult males have revealed a decrease in abundance during the first 8 weeks of adulthood (17, 45), possibly suggesting that the degenerative effect of host ageing on the symbiosis occurs sooner in male hosts than their female counterparts.

We proposed that senescence of the immune system may allow the population of *Sodalis* to increase again in very old flies (Fig 1a), which we observed in females from the age of approximately 67 days old (Fig 2b). In contrast to obligate symbionts, little is known of the mechanisms by which hosts control the abundance of facultative symbionts. Studies on the *Riptortus* bean bug have revealed that the host uses antimicrobial peptides (AMPs) to regulate the abundance of the facultative *Burkholderia* (63), however, the interactions between *Sodalis* and the tsetse immune system appear complex. Trappeniers *et al*. (2019) reported a reduced immune response to *Sodalis* (64), potentially allowing its colonisation throughout the host lifetime, however, Weiss *et al*. (2008) reported that recognition of *Sodalis* surface proteins by the host induce the expression of immunity related genes (65). Additionally, *Sodalis* have demonstrated resistance to multiple host AMPs (66-68), and this symbiont may be able to evade host immune effectors by invading host cells (69). It is likely that the immune system is finely tuned to the presence of *Sodalis* to limit its proliferation without removing it completely (70). An alternative explanation is that *Sodalis* growth is self-regulated. *Sodalis* has demonstrated the ability to suppress virulence in tsetse via quorum sensing (71), which potentially limits the burden of this symbiont on its host. It might be beneficial for the symbiont to limit its own proliferation during early adulthood to allow host investment in reproduction, as the fitness interests of *Sodalis* and the host are aligned whilst the host is reproducing due to vertical symbiont transmission (72). However, exploitation of host resources and unregulated symbiont growth could increase *Sodalis* fitness in hosts that are no longer reproducing as increased symbiont density may promote the likelihood of horizontal transmission. Indeed, laboratory-reared female tsetse experience reproductive senescence, whereby the likelihood of larval abortion increases and provisioning of resources to offspring decreases with increasing maternal age (73, 74). The mechanism by which *Sodalis* invades host cells indicates that it evolved from parasitic ancestors (75), and it is plausible that a facultative symbiont, which has recently transitioned to host-associated living, may express selfish and opportunistic traits characteristic of parasitism under certain contexts.

We observed that the density of *Wigglesworthia* genomes is generally greater than that of *Sodalis* within the digestive and reproductive tissues (Fig 2). This was consistent among our three studies, which tested both males and females of a range of ages (Figs 2,3), and is consistent with previous studies which measured symbiont densities within whole flies (17, 45). It is potentially surprising given that *Sodalis* demonstrates a wider tissue tropism than *Wigglesworthia* and does not rely on the host to mediate cell replication (35). As our primer pairs were designed to target single-copy genes in both symbionts and tsetse, it is possible that *Wigglesworthia* demonstrates a high prevalence of polyploidy which *Sodalis* does not, or that the host can direct a large amount of resources into the bacteriome for maintenance of *Wigglesworthia*.

Proliferation of midgut bacteria following a blood meal is demonstrated in other blood-feeding insects, such as mosquitoes, peaking at 24-36 hours post-blood feed (49). We hypothesised that *Sodalis* would demonstrate a comparable proliferation, due to its presence in the midgut, however we found no evidence of changes in the density of either *Wigglesworthia* or *Sodalis* throughout the hunger cycle (Fig 2c,d). The gut symbionts of adult mosquitoes are acquired environmentally, and although *Sodalis* is a facultative symbiont we consider that there is still an evolutionary history which may subject *Sodalis* to a greater degree of regulation – whether host- or symbiont-mediated – than transient symbionts of mosquitoes. The growth of cultured *Sodalis* has been shown to be slow relative to other bacteria, with a doubling time of approximately 26 hours (76). The growth kinetics *in vitro* may correspond to the within-host growth of *Sodalis*, in which case density fluctuation within a hunger cycle is likely to be weak. It may also be that we did not have the power to detect small, within-individual effects if they existed due to our study design, where measurements were taken from different hosts (by necessity), and large variation in symbiont density may exist between individuals.

By rearing tsetse on different diets, we aimed to manipulate the requirements of hosts for symbiont-derived micronutrients. We hypothesised that hosts would regulate the abundance of *Wigglesworthia* to maximise the benefit of hosting this symbiont population, whereas the growth of *Sodalis* would respond directly to the availability of nutrients in the diet (Fig 1c). We did not find strong evidence of the density of *Wigglesworthia* changing in hosts reared on different diets (Fig 3a,b). We consider two explanations. First, it is possible that the cost of maintaining *Wigglesworthia* is relatively low, such that even on a diet with very little macronutrient content, there is still a net benefit to hosting a large *Wigglesworthia* density. Second, the cost of reducing *Wigglesworthia* may be substantial, and it may not be beneficial to actively regulate according to host nutrition. The intracellular localisation of *Wigglesworthia* support the second suggestion, as reduction in the cell density of *Wigglesworthia* may require autophagy of the host’s own bacteriocytes (16, 61, 70). Aphids are reported to harbour different densities of their obligate symbiont (which resides within their bacteriome) according to their host plant (13). In contrast to aphids which feed on one plant, however, tsetse do not feed from one host individual but find a new blood meal every few days. Tsetse may therefore regulate *Wigglesworthia* according to long-term nutritional requirements rather than short-term fluctuations in diet. The stationary dynamics of *Wigglesworthia* density observed throughout the hunger cycle (Fig 2c) provides additional support to the fact that tsetse may maintain a constant, optimal density of *Wigglesworthia*. When the relative cost of supporting *Wigglesworthia* is great (such as when dietary energy is limited), or where the requirements for *Wigglesworthia* are reduced (such as when the host receives a higher quality diet) it is possible that tsetse reduce *Wigglesworthia* metabolism as an alternative to reducing cell density (11, 77).

In contrast to *Wigglesworthia*, we found that the density of *Sodalis* changed with dietary manipulation as predicted (Fig 1c), whereby the density of *Sodalis* was lower in hosts reared on diets with lower blood content (Fig 3c), and greater in hosts reared on diets enriched with yeast, an exogenous source of B vitamins (Fig 3d). Reduced macronutrient availability, as well as decreased thiamine production by *Wigglesworthia* (38), in tsetse receiving diets of diluted blood may limit the growth of *Sodalis*. As there is no clear benefit of hosting a larger population of *Sodalis* when tsetse are supplied with a higher quality diet, we infer that the resulting increase in *Sodalis* density in hosts receiving blood enriched with yeast extract is not the result of tsetse regulating *Sodalis* to a higher density, rather that this is mediated by *Sodalis* itself. Rearing tsetse on a diet of diluted blood imposed detrimental effects on host reproduction, indicated by the reduced pupae weight and starvation time of offspring upon emergence (Fig 4a,c), which is consistent with previous findings (73). Enriching blood with yeast extract also impacted reproduction (Fig 4b,d) and severely reduced survival in hosts administered 5% yeast extract (Fig S1), possibly indicating a toxic effect of this treatment. A reduction in tsetse fecundity has previously been observed with supplementation of host diet with yeast extract (50). Hosts experiencing nutritional stress possibly also suffer from impaired immunity (78). As well as being provided with a greater complement of nutrients, the host may have a reduced ability to limit *Sodalis* proliferation, hence the increased density of *Sodalis* (Fig 4d). Any detrimental effects on immunity caused by the dietary treatments, however, do not appear to affect the density of *Wigglesworthia* (Fig 3a,b).

## Conclusion

We found evidence that the ecological contexts that influence symbiont growth and density in tsetse are different for *Wigglesworthia* and *Sodalis*, their obligate and facultative symbionts respectively. Although we observed a decline in *Wigglesworthia* density in older flies, we found no evidence that *Wigglesworthia* density is actively adjusted to meet short-term nutritional requirements; rather we suggest that it is maintained at a high abundance by the host to maximise long-term benefits. Variation in nutrient availability appears to have little impact on the density of *Wigglesworthia*, supporting the hypothesis that density is primarily controlled by the host. *Sodalis*, however, appears to respond dynamically to the conditions of its environment, and factors such as host age and nutrient availability have significant impacts on the density of this symbiont. We suggest that due to the recent transition to host-associated living, both the host and symbiont possess a degree of control over the density of this facultative symbiont, potentially via host immune function and self-regulation. The potential for vertically transmitted, facultative symbionts to transition along the mutualism-parasitism spectrum according to the ecological context, and the evolutionary outcomes of such interactions, warrant further investigation.

## Acknowledgments

The authors thank Dr Hester Weaving and Dr Lori Peacock for providing tsetse husbandry training to M. Whittle; IAEA (Vienna, Austria) for supplying *G. morsitans* pupae for practice dissections and qPCR optimisation; Avi Leong and Kyle Du for assisting with measuring tsetse wings; Dr Drew Allen and Shannon Kaiser for statistics advice; and Prof. Richard Wall and Dr Ajay Narendra for use of equipment and laboratory space.

## Funding

This work was supported by an EPSRC/University of Bristol/Macquarie University Cotutelle studentship to M. Whittle (EP/T517872/1) and Biotechnology and Biological Sciences Research Council grant to S. English and A.M.G. Barreaux (BB/P006159/1). S. English is also supported by a Royal Society Dorothy Hodgkin Fellowship (DH140236) and UK Research and Innovation Future Leaders Fellowship (MR/W007711/1).

## Notes

### Competing Interest Statement

The authors have declared no competing interest.

## References

1. Buchner P. 1965 Endosymbiosis of animals with plant microorganisms. New York, USA: Interscience publishers.

2. Husnik F, Tashyreva D, Boscaro V, George EE, Lukes J, Keeling PJ. 2021 Bacterial and archaeal symbioses with protists. Curr. Biol. 31, R862–R77.

3. Muller DB, Vogel C, Bai Y, Vorholt JA. 2016 The plant microbiota: systems-level insights and perspectives. Annu. Rev. Genet. 50, 211–34.

4. Drew GC, Stevens EJ, King KC. 2021 Microbial evolution and transitions along the parasite-mutualist continuum. Nat. Rev. Microbiol. 19, 623–38.

5. Venn AA, Loram JE, Douglas AE. 2008 Photosynthetic symbioses in animals. J. Exp. Bot. 59, 1069–80.

6. Flórez L, Biedermann PHW, Engl T, Kaltenpoth M. 2015 Defensive symbioses of animals with prokaryotic and eukaryotic microorganisms. Nat. Prod. Rep. 32, 904–36.

7. Hector TE, Hoang KL, Li J, King KC. 2022 Symbiosis and host responses to heating. Trends Ecol. Evol. 37, 611–24.

8. Chomicki G, Kiers ET, Renner SS. 2020 The evolution of mutualistic dependence. Annu. Rev. Ecol. Evol. Syst. 51, 409–32.

9. Ankrah NYD, Chouaia B, Douglas AE. 2018 The cost of metabolic interactions in symbioses between insects and bacteria with reduced genomes. mBio 9, e01433–18.

10. Simonet P, Duport G, Gaget K, Weiss-Gayet M, Colella S, Febvay G, et al. 2016 Direct flow cytometry measurements reveal a fine-tuning of symbiotic cell dynamics according to the host developmental needs in aphid symbiosis. Sci. Rep. 6, 19967.

11. Snyder AK, McLain C, Rio RV. 2012 The tsetse fly obligate mutualist Wigglesworthia morsitans alters gene expression and population density via exogenous nutrient provisioning. Appl. Environ. Microbiol. 78, 7792–7.

12. Wilkinson TL, Koga R, Fukatsu T. 2007 Role of host nutrition in symbiont regulation: impact of dietary nitrogen on proliferation of obligate and facultative bacterial endosymbionts of the pea aphid Acyrthosiphon pisum. Appl. Environ. Microbiol. 73, 1362–6.

13. Zhang YC, Cao WJ, Zhong LR, Godfray HCJ, Liu XD. 2016 Host plant determines the population size of an obligate symbiont (Buchnera aphidicola) in aphids. Appl. Environ. Microbiol. 82, 2336–46.

14. Scarborough CL, Ferrari J, Godfray HC. 2005 Aphid protected from pathogen by endosymbiont. Science 310, 1781.

15. Russell JA, Moran NA. 2006 Costs and benefits of symbiont infection in aphids: variation among symbionts and across temperatures. Proc. Biol. Sci. 273, 603–10.

16. Vigneron A, Masson F, Vallier A, Balmand S, Rey M, Vincent-Monegat C, et al. 2014 Insects recycle endosymbionts when the benefit is over. Curr. Biol. 24, 2267–73.

17. Rio RV, Wu YN, Filardo G, Aksoy S. 2006 Dynamics of multiple symbiont density regulation during host development: tsetse fly and its microbial flora. Proc. Biol. Sci. 273, 805–14.

18. Serrato-Salas J, Gendrin M. 2023 Involvement of microbiota in insect physiology: focus on B vitamins. mBio 14, e02225–22.

19. Douglas AE. 1989 Mycetocyte symbiosis in insects. Biol. Rev. Camb. Philos. Soc. 64, 409–34.

20. Holland JN, DeAngelis DL, Schultz ST. 2004 Evolutionary stability of mutualism: interspecific population regulation as an evolutionarily stable strategy. Proc. Biol. Sci. 271, 1807–14.

21. Whittle M, Barreaux AMG, Bonsall MB, Ponton F, English S. 2021 Insect-host control of obligate, intracellular symbiont density. Proc. Biol. Sci. 288, 20211993.

22. Haines LR, Vale GA, Barreaux AMG, Ellstrand NC, Hargrove JW, English S. 2020 Big baby, little mother: tsetse flies are exceptions to the juvenile small size principle. Bioessays 42, e2000049.

23. Nogge G, Ritz R. 1982 Number of symbionts and its regulation in tsetse flies, Glossina spp. Entomol. Exp. Appl. 31, 249–54.

24. Husnik F, Keeling PJ. 2019 The fate of obligate endosymbionts: reduction, integration, or extinction. Curr. Opin. Genet. Dev. 58-59, 1–8.

25. McCutcheon JP, Moran NA. 2011 Extreme genome reduction in symbiotic bacteria. Nat. Rev. Microbiol. 10, 13–26.

26. Chomicki G, Werner GDA, West SA, Kiers ET. 2020 Compartmentalization drives the evolution of symbiotic cooperation. Philos. Trans. R. Soc. Lond. B Biol. Sci. 375, 20190602.

27. Nguyen PL, van Baalen M. 2020 On the difficult evolutionary transition from the free-living lifestyle to obligate symbiosis. PLoS One 15, e0235811.

28. Kikuchi Y. 2009 Endosymbiotic bacteria in insects: their diversity and culturability. Microbes Environ. 24, 195–204.

29. Wang J, Weiss BL, Aksoy S. 2013 Tsetse fly microbiota: form and function. Front. Cell. Infect. Microbiol. 3, 69.

30. Chen X, Li S, Aksoy S. 1999 Concordant evolution of a symbiont with its host insect species: molecular phylogeny of genus Glossina and its bacteriome-associated endosymbiont, Wigglesworthia glossinidia. J. Mol. Evol. 48, 49–58.

31. Moran NA, Bennett GM. 2014 The tiniest tiny genomes. Annu. Rev. Microbiol. 68, 195–215.

32. Akman L, Aksoy S. 2001 A novel application of gene arrays: Escherichia coli array provides insight into the biology of the obligate endosymbiont of tsetse flies. Proc. Natl. Acad. Sci. 98, 7546–51.

33. Nogge G. 1976 Sterility in tsetse flies (Glossina morsitans Westwood) caused by loss of symbionts. Experientia 32, 995–6.

34. Benoit JB, Attardo GM, Baumann AA, Michalkova V, Aksoy S. 2015 Adenotrophic viviparity in tsetse flies: potential for population control and as an insect model for lactation. Annu. Rev. Entomol. 60, 351–71.

35. Balmand S, Lohs C, Aksoy S, Heddi A. 2013 Tissue distribution and transmission routes for the tsetse fly endosymbionts. J. Invertebr. Pathol. 112, S116–22.

36. Toh H, Weiss BL, Perkin SA, Yamashita A, Oshima K, Hattori M, et al. 2006 Massive genome erosion and functional adaptations provide insights into the symbiotic lifestyle of Sodalis glossinidius in the tsetse host. Genome Res. 16, 149–56.

37. De Vooght L, Caljon G, Van Hees J, Van Den Abbeele J. 2015 Paternal transmission of a secondary symbiont during mating in the viviparous tsetse fly. Mol. Biol. Evol. 32, 1977–80.

38. Snyder AK, Deberry JW, Runyen-Janecky L, Rio RV. 2010 Nutrient provisioning facilitates homeostasis between tsetse fly (Diptera: Glossinidae) symbionts. Proc. Biol. Sci. 277, 2389–97.

39. Welburn SC, Maudlin I, Ellis DS. 1987 In vitro cultivation of rickettsia-like-organisms from Glossina spp. Ann. Trop. Med. Parasitol. 81, 331–5.

40. Dale C, Welburn SC. 2001 The endosymbionts of tsetse flies: manipulating host-parasite interactions. Int. J. Parasitol. 31, 628–31.

41. Dossi FC, da Silva EP, Consoli FL. 2014 Population dynamics and growth rates of endosymbionts during Diaphorina citri (Hemiptera, Liviidae) ontogeny. Microb. Ecol. 68, 881–9.

42. Wolschin F, Holldobler B, Gross R, Zientz E. 2004 Replication of the endosymbiotic bacterium Blochmannia floridanus is correlated with the developmental and reproductive stages of its ant host. Appl. Environ. Microbiol. 70, 4096–102.

43. Stoll S, Feldhaar H, Gross R. 2009 Transcriptional profiling of the endosymbiont Blochmannia floridanus during different developmental stages of its holometabolous ant host. Environ. Microbiol. 11, 877–88.

44. Stoll S, Feldhaar H, Fraunholz MJ, Gross R. 2010 Bacteriocyte dynamics during development of a holometabolous insect, the carpenter ant Camponotus floridanus. BMC Microbiol. 10, 308.

45. Hamidou Soumana I, Berthier D, Tchicaya B, Thevenon S, Njiokou F, Cuny G, et al. 2013 Population dynamics of Glossina palpalis gambiensis symbionts, Sodalis glossinidius, and Wigglesworthia glossinidia, throughout host-fly development. Infect. Genet. Evol. 13, 41–8.

46. Haines LR. 2013 Examining the tsetse teneral phenomenon and permissiveness to trypanosome infection. Front. Cell. Infect. Microbiol. 3, 84.

47. Hillyer JF. 2016 Insect immunology and hematopoiesis. Dev. Comp. Immunol. 58, 102–18.

48. Gusmao DS, Santos AV, Marini DC, Bacci M, Jr., Berbert-Molina MA, Lemos FJ. 2010 Culture-dependent and culture-independent characterization of microorganisms associated with Aedes aegypti (Diptera: Culicidae) (L.) and dynamics of bacterial colonization in the midgut. Acta Trop. 115, 275–81.

49. Oliveira JH, Goncalves RL, Lara FA, Dias FA, Gandara AC, Menna-Barreto RF, et al. 2011 Blood meal-derived heme decreases ROS levels in the midgut of Aedes aegypti and allows proliferation of intestinal microbiota. PLoS Pathog. 7, e1001320.

50. Alam U, Medlock J, Brelsfoard C, Pais R, Lohs C, Balmand S, et al. 2011 Wolbachia symbiont infections induce strong cytoplasmic incompatibility in the tsetse fly Glossina morsitans. PLoS Pathog. 7, e1002415.

51. Whittle M, Bonsall MB, Barreaux AMG, Ponton F, English S. 2023 A theoretical model for host-controlled regulation of symbiont density. J. Evol. Biol. 36, 1731–44.

52. Hargrove J, English S, Torr SJ, Lord J, Haines LR, van Schalkwyk C, et al. 2019 Wing length and host location in tsetse (Glossina spp.): implications for control using stationary baits. Parasit. Vectors 12, 24.

53. Langley PA. 1977 Physiology of tsetse flies (Glossina spp.) (Diptera: Glossinidae): a review. Bull. Entomol. Res. 67, 523–74.

54. Team RC. R Foundation for statistical computing. 2022 R: A language and environment for statistical computing. Vienna, Austria

55. Wagenmakers E-J, Farrel S. 2004 AIC model selection using Akaike weights. Psych. Bull. Rev. 11, 192–6.

56. Burnham KP, Anderson DR. 2002 Model selection and multimodel inference: a practical information-theoretic approach. New York, USA: Springer.

57. Lüdecke D, Ben-Shachar MS. 2021 performance: an R package for assessment, comparison and testing of statistical models. J. Open Source Softw. 6, 3139.

58. Jackson CH. 1949 The biology of tsetse flies. Biol. Rev. Camb. Philos. Soc. 24, 174–99.

59. Leak SG. 1998 Tsetse biology and ecology: their role in the epidemiology and control of trypanosomiasis. Oxfordshire, UK: CABI Publishing.

60. Ferrarini MG, Dell’Aglio E, Vallier A, Balmand S, Vincent-Monegat C, Hughes S, et al. 2022 Efficient compartmentalization in insect bacteriomes protects symbiotic bacteria from host immune system. Microbiome 10, 156.

61. Simonet P, Gaget K, Balmand S, Ribeiro Lopes M, Parisot N, Buhler K, et al. 2018 Bacteriocyte cell death in the pea aphid/Buchnera symbiotic system. Proc. Natl. Acad. Sci. U.S.A. 115, E1819–E28.

62. Komaki K, Ishikawa H. 2000 Genomic copy number of intracellular bacterial symbionts of aphids varies in response to developmental stage and morph of their host. Insect Biochem. Mol. Biol. 30, 253–8.

63. Kim JK, Lee JB, Jang HA, Han YS, Fukatsu T, Lee BL. 2016 Understanding regulation of the host-mediated gut symbiont population and the symbiont-mediated host immunity in the Riptortus-Burkholderia symbiosis system. Dev. Comp. Immunol. 64, 75–81.

64. Trappeniers K, Matetovici I, Van Den Abbeele J, De Vooght L. 2019 The tsetse fly displays an attenuated immune response to its secondary symbiont, Sodalis glossinidius. Front. Microbiol. 10, 1650.

65. Weiss BL, Wu Y, Schwank JJ, Tolwinski NS, Aksoy S. 2008 An insect symbiosis is influenced by bacterium-specific polymorphisms in outer-membrane protein A. Proc. Natl. Acad. Sci. U.S.A. 105, 15088–93.

66. Hu Y, Aksoy S. 2005 An antimicrobial peptide with trypanocidal activity characterized from Glossina morsitans morsitans. Insect Biochem. Mol. Biol. 35, 105–15.

67. Hao Z, Kasumba I, Lehane MJ, Gibson WC, Kwon J, Aksoy S. 2001 Tsetse immune responses and trypanosome transmission: Implications for the development of tsetse-based strategies to reduce trypanosomiasis. Proc. Natl. Acad. Sci. 98, 12648–53.

68. Haines LR, Hancock REW, Pearson TW. 2003 Cationic antimicrobial peptide killing of African trypanosomes and Sodalis glossinidius, a bacterial symbiont of the insect vector of sleeping sickness. Vector-borne and Zoonotic diseases 3, 175–86.

69. Thakur A, Mikkelsen H, Jungersen G. 2019 Intracellular pathogens: host immunity and microbial persistence strategies. J. Immunol. Res. 1, 1356540.

70. Ratzka C, Gross R, Feldhaar H. 2012 Endosymbiont tolerance and control within insect hosts. Insects 3, 553–72.

71. Medina Munoz M, Spencer N, Enomoto S, Dale C, Rio RVM. 2020 Quorum sensing sets the stage for the establishment and vertical transmission of Sodalis praecaptivus in tsetse flies. PLoS Genet. 16, e1008992.

72. Mushegian AA, Ebert D. 2016 Rethinking “mutualism” in diverse host-symbiont communities. Bioessays 38, 100–8.

73. Lord JS, Leyland R, Haines LR, Barreaux AMG, Bonsall MB, Torr SJ, et al. 2021 Effects of maternal age and stress on offspring quality in a viviparous fly. Ecol. Lett. 24, 2113–22.

74. Barreaux AMG, Higginson AD, Bonsall MB, English S. 2022 Incorporating effects of age on energy dynamics predicts nonlinear maternal allocation patterns in iteroparous animals. Proc. Biol. Sci. 289, 20211884.

75. Dale C. 2001 From the Cover: The insect endosymbiont Sodalis glossinidius utilizes a type III secretion system for cell invasion. Proc. Natl. Acad. Sci. 98, 1883–8.

76. Matthew CZ, Darby AC, Young SA, Hume LH, Welburn SC. 2005 The rapid isolation and growth dynamics of the tsetse symbiont Sodalis glossinidius. FEMS Microbiol. Lett. 248, 69–74.

77. Smith TE, Moran NA. 2020 Coordination of host and symbiont gene expression reveals a metabolic tug-of-war between aphids and Buchnera. Proc. Natl. Acad. Sci. U.S.A. 117, 2113–21.

78. Ponton F, Tan YX, Forster CC, Austin AJ, English S, Cotter SC, et al. 2023 The complex interactions between nutrition, immunity and infection in insects. J. Exp. Biol. 226, 24.

